# Equi-depth pooling for standardized and cost-effective Adaptive Immune Receptor Repertoire sequencing (AIRRseq)

**DOI:** 10.64898/2026.07.29.740903

**Authors:** T.W. Verdonckt, M. Sahulčík, C. Struyfs, A.S. Vermeersch, K. Bartholomeeusen, F. Van Nieuwerburgh, O. Lagatie, K. K. Ariën

## Abstract

Bulk B- and T-cell adaptive immune receptor (BCR/TCR) repertoire sequencing (AIRRseq) enables comprehensive analysis of adaptive immune diversity, but achieving balanced sequencing depth across heterogeneous clinical samples remains a major cost driver. Standard equimolar pooling prior to sequencing disproportionately allocates reads, leading to undersequencing of libraries with high sequence abundance and oversampling of those with few sequences.

Here, a quantitative pooling strategy is described in which libraries are pooled based on estimated target-specific unique molecular identifier (UMI) counts rather than library molarity, ensuring a uniform number of supporting reads per target sequence UMI across samples. Libraries were prepared from human RNA using the NEBNext Immune Sequencing (IS) kit and quantified by qPCR during the second amplification (PCR2) step. Deeply sequenced samples from a dengue 1 human infection model (DHIM1) clinical study were used to model the log-linear relationship between PCR2 cycle threshold (Ct) values and detected sequence counts. This model was then applied to a second, independent batch of samples from a dengue 3 human infection model (DHIM3) clinical study to predict UMI abundance and guide equi-depth pooling prior to sequencing.

Sequencing of these equi-depth pools demonstrated a near-uniform reads-per-sequence ratio across samples, confirming that this approach achieves balanced depth without oversampling and at reduced cost.

Simulations across a range of between-sample UMI-count distributions reflecting real-world sample pools showed that equi-depth pooling is expected to reduce the total number of required sequencing reads by approximately 70% compared with equimolar pooling, with savings increasing as the spread of library sizes grows. Beyond cost efficiency, equi-depth pooling eliminates the complexity-dependent sampling bias inherent to equimolar pooling, ensuring uniform sequencing depth and comparable UMI recovery across small and large libraries. For new workflows or sample types, implementation requires either an existing calibration dataset or a pilot sequencing experiment to establish the relationship between PCR2 Ct values and relative UMI abundance.

The equi-depth method thus provides a robust, scalable, and cost-efficient strategy for bulk AIRRseq studies where library sizes vary widely.

## Introduction

Bulk sequencing of B- and T-cell receptor (BCR/TCR) repertoires enables quantitative characterization of clonal diversity and dynamics across large cohorts at a fraction of the cost of single-cell approaches, and with substantially greater coverage per sample. Reviews and guidelines emphasize that key experimental choices (including starting material, library construction, UMI usage, and sequencing depth) critically influence quantitative accuracy^1–5^.

Among bulk BCR/TCR sequencing strategies, 5′ rapid amplification of cDNA ends (5′ RACE) performed on total RNA using primers targeting the constant region is generally regarded as good practice because it captures the full V(D)J–C transcript without requiring prior knowledge of variable region germline alleles and thereby avoids primer-annealing biases inherent to multiplex PCR schemes. In addition, RNA-based 5′ RACE allows identification of immunoglobulin isotypes and productive transcripts while maintaining quantitative representation when coupled to unique molecular identifiers (UMIs) for molecule counting and consensus assembly. Consensus assembly mitigates amplification/sequencing errors to support accurate clone counting and lineage analysis^6^.

Adequate sequencing depth is central to robust repertoire analyses^7^. Even very deep profiling may not exhaustively recover clonotypes as shown by rarefaction/species-richness analyses of highly sampled human repertoires evidenced by asymptotically increasing but non-plateauing curves, indicating persistent unseen diversity despite extreme read counts. This phenomenon has been documented explicitly in deeply sequenced human BCR datasets underscoring the need to plan sequencing depth with unseen clonotype mass in mind, deploy estimators such as Chao1 when appropriate, and minimize undersampling-induced biases^8^.

Because somatic hypermutation introduces true nucleotide-level variation that can resemble PCR or sequencing errors, UMI-derived immune receptor sequences require sufficient read support for reliable consensus construction. In the nf-core/airrflow workflow used here, sequences supported by fewer than two representative reads are filtered out by default, so effective repertoire recovery depends on achieving adequate reads per UMI^9^. Consequently, the required number of reads is directly determined by the number of unique sequences in a sample. Samples with higher diversity therefore yield fewer usable sequences than smaller samples, even when allocated more raw reads, if only a fraction of sequences receive more than one supporting read. This emphasizes that sequencing depth planning must consider both total sequence load as well as the distribution of reads across sequences to avoid loss of information in high-complexity samples^10^.

Earlier methodological work quantifying detection power shows that the probability of detecting clones depends strongly on sequencing depth relative to library size and clone frequency, reinforcing that depth allocation must account for library-specific molecule counts to avoid technological undersampling biases^11^.

This mismatch between sequencing depth and molecular complexity has also been noted in studies quantifying UMI-based library efficiency, where a large proportion of reads collapse to a small number of unique molecules if complexity is low. For instance, bioinformatic analysis of liquid biopsy libraries with UMI deduplication showed that effective unique coverage was reduced by ∼60-fold relative to raw read depth, highlighting the inefficiency of uniform read allocation when molecule counts differ between libraries^12^.

Clinical contexts further complicate sequencing depth planning because library sizes vary widely across samples. For example, acute viral infections such as dengue can drive >100-fold plasmablast expansions in peripheral blood^11^, producing comparable orders-of-magnitude differences in RNA template and UMI yield across time points and individuals. A common operational choice relies on equimolar pooling of libraries, allocating equal read counts to the sequencing libraries regardless of UMI abundance per library. In pools of heterogeneous size, this practice will result in varying read/UMI counts. If a lower threshold read/UMI count *r* is desired for all samples, the total required read count is defined by

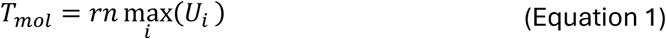

where *n* is the number of samples, *U_i_* the expected UMI count for sample *i* and *T_mol_* the total raw reads required. In heterogeneous pools, this makes the total sequencing requirement scale with the most UMI-rich library, which can render sequencing runs cost-prohibitive. If this threshold is not reached, UMI-rich libraries are systematically undersequenced, reducing read support per molecule and biasing repertoire recovery toward higher-abundance clonotypes in the most complex samples.

While ad hoc adjustments and separate sequencing runs are sometimes used to handle high-variance libraries, this introduces additional work and time cost. To date, systematic pooling strategies based on molecular counts remain uncommon. Outside AIRRseq, related ideas have been applied in contexts like whole-genome sequencing of organisms with different genome sizes, where coverage-based pooling is used to avoid over- or under-sequencing^13^. The same principle applies to UMI-based RNA libraries, where allocating reads based on estimated molecule count can equalize information yield across libraries.

The equi-depth method presented here is tailored for RNA-based 5′ RACE + UMI workflows, like that used by the NEBNext® Immune Sequencing Kit (Human), as it relies on qPCR steps that are already performed in standard library preparation (Figure 1). In such protocols, cDNA is obtained through template-switch reverse transcription with UMIs. The cDNA is then amplified with target-specific primers during the ‘PCR1’ step with 12 amplification cycles, after which adapters are added and the libraries amplified to a desired amount using sample-specific cycle counts during the ‘PCR2’ step. To determine the number of cycles required for each sample in PCR2, an optional qPCR step is performed, where the cycle threshold is set as two-thirds of the maximal fluorescence (*Ct*_2/3_)^14^.

**Figure 1 –.**
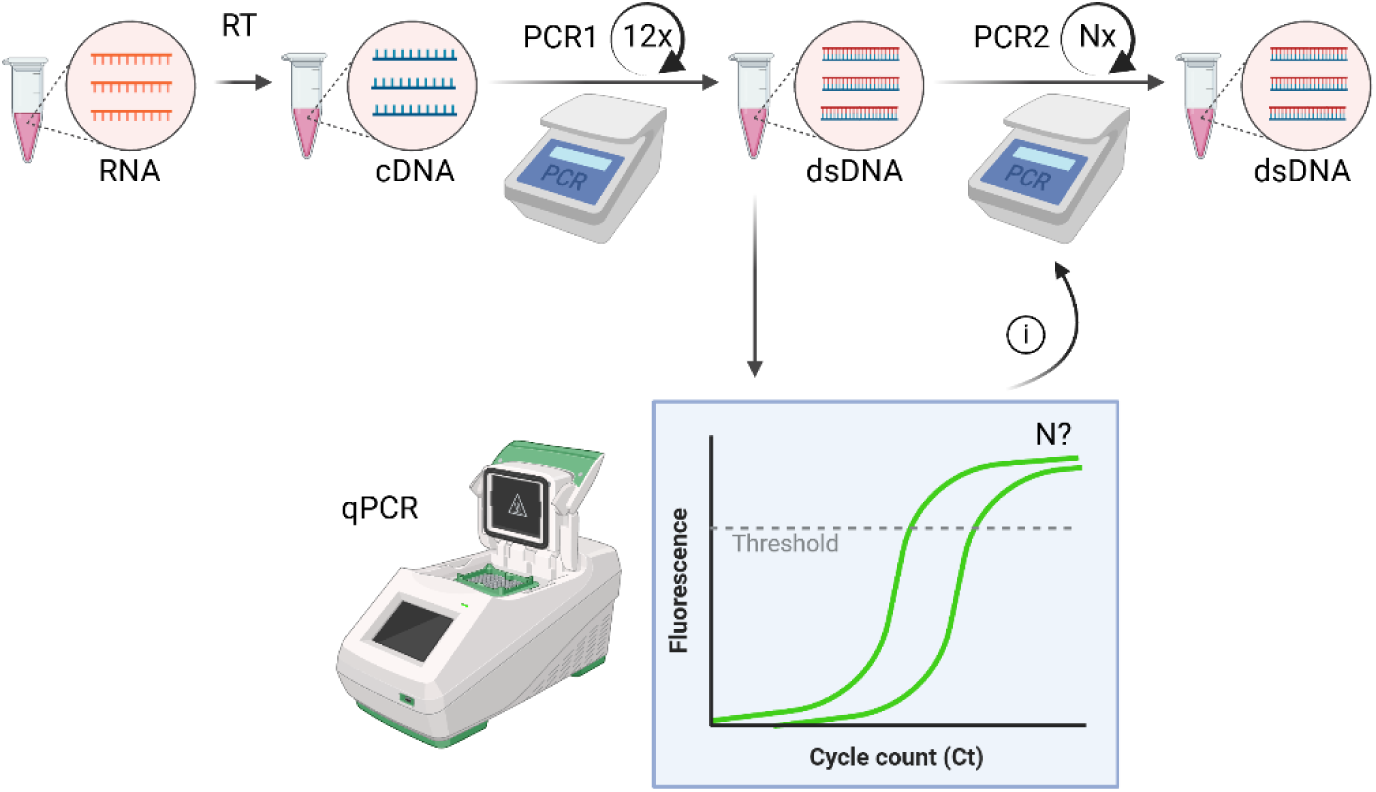
NEBNext® Immune Sequencing Kit workflow. First, cDNA is obtained through reverse transcription with template-switch oligos containing UMI sequences. The cDNA is then used as the template in target-specific PCR reaction (PCR1) that is run for 12 cycles. Hereafter, a qPCR step is performed to assess the number of cycles required to further amplify the DNA till two-thirds of the maximal fluorescence. This cycle threshold value is then used in the PCR2 reaction which yields ready-to-sequence libraries within the desired amount range.

In the current work, we hypothesized that equimolar pooling is suboptimal for large, diverse sample pools because it does not equalize reads per UMI. A quantitative pooling strategy was therefore devised in which pool fractions are set proportional to estimated UMI counts (rather than library molarity) so that each library receives approximately the same reads/UMI at sequencing. The total read count to achieve a desired read/UMI threshold *r* would then be defined as

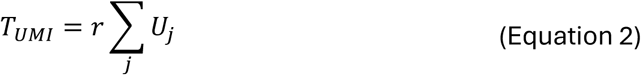

which results in a strong total read count reduction compared to equimolar pooling (Equation 1). To realize this in practice, the log-linear relationship between the count of cycles required to reach 2/3^rd^ of maximal fluorescence in the PCR2 step (*Ct*_2/3_) and the count of underlying UMIs in the resulting sequencing library was modelled using deeply sequenced libraries to provide a calibration curve; this calibration was subsequently applied to batches from an independent clinical study of dengue human infection to set pooling ratios prospectively. The conceptual motivation aligns with prior recommendations on study design and sampling/coverage considerations in AIRRseq experiments and with statistical work on richness estimation and under-sampling^2^.

This study (i) establishes the Ct→UMI model from analysis of oversequenced libraries; (ii) applies the model to predict UMI abundance from a qPCR step for a second clinical batch, to (iii) perform equi-depth pooling; and (iv) evaluates technical outcomes (uniformity of reads/UMI and cost savings) in real data and simulations.

Simulated pools spanning increasing skewness, defined as progressively stronger right-tailed UMI-count distributions in which a minority of libraries contains a disproportionate fraction of total UMIs, demonstrate that equi-depth pooling reduces total required reads compared with equimolar pooling. The savings increase with between-sample UMI-count heterogeneity, and real clinical cohorts with highly unequal library sizes are therefore expected to benefit accordingly.

Importantly, the approach described here is not restricted to this specific chemistry and can be applied to other library-preparation methods and sequencing platforms, provided that the workflow includes a PCR amplification step in which libraries are amplified to a defined target molarity or yield.

## Results

### Calibration of qPCR cycle threshold (*Ct*_2/3_) to UMI count

To enable equi-depth pooling, the relationship between PCR2 amplification kinetics of the library-preparation protocol (*Ct*_2/3_) and representative UMI abundance was first established by performing deep sequencing of DHIM1^15^ study samples. DHIM1 RNA samples were used to generate parallel bulk TCR and IGH libraries using the NEBNext Immune Sequencing workflow as described in STAR★METHODS. The prepared libraries were quantified and pooled equimolarly to a concentration of 10 µM; the pool sequenced and the raw reads assembled with the nf-core/airrflow analysis pipeline^9^ version 5.0.0 (STAR★METHODS). Sequencing depth was assessed separately for each library and target locus by plotting the read-per-UMI distribution of annotated sequences represented by a single UMI (Figure 2). Libraries were considered sufficiently sequenced for Ct–UMI calibration when the modal read-per-UMI value for the retained target loci was approximately 10 or higher. This empirical criterion was chosen because, at this depth, the lower tail of the read-per-UMI distribution remains largely above the two-read support threshold used during repertoire assembly, minimizing loss of UMI-derived sequences due to insufficient read support. Libraries for which one or more target loci showed modal read support at or below this range were considered at risk of UMI undercounting and were excluded from calibration.

**Figure 2 –.**
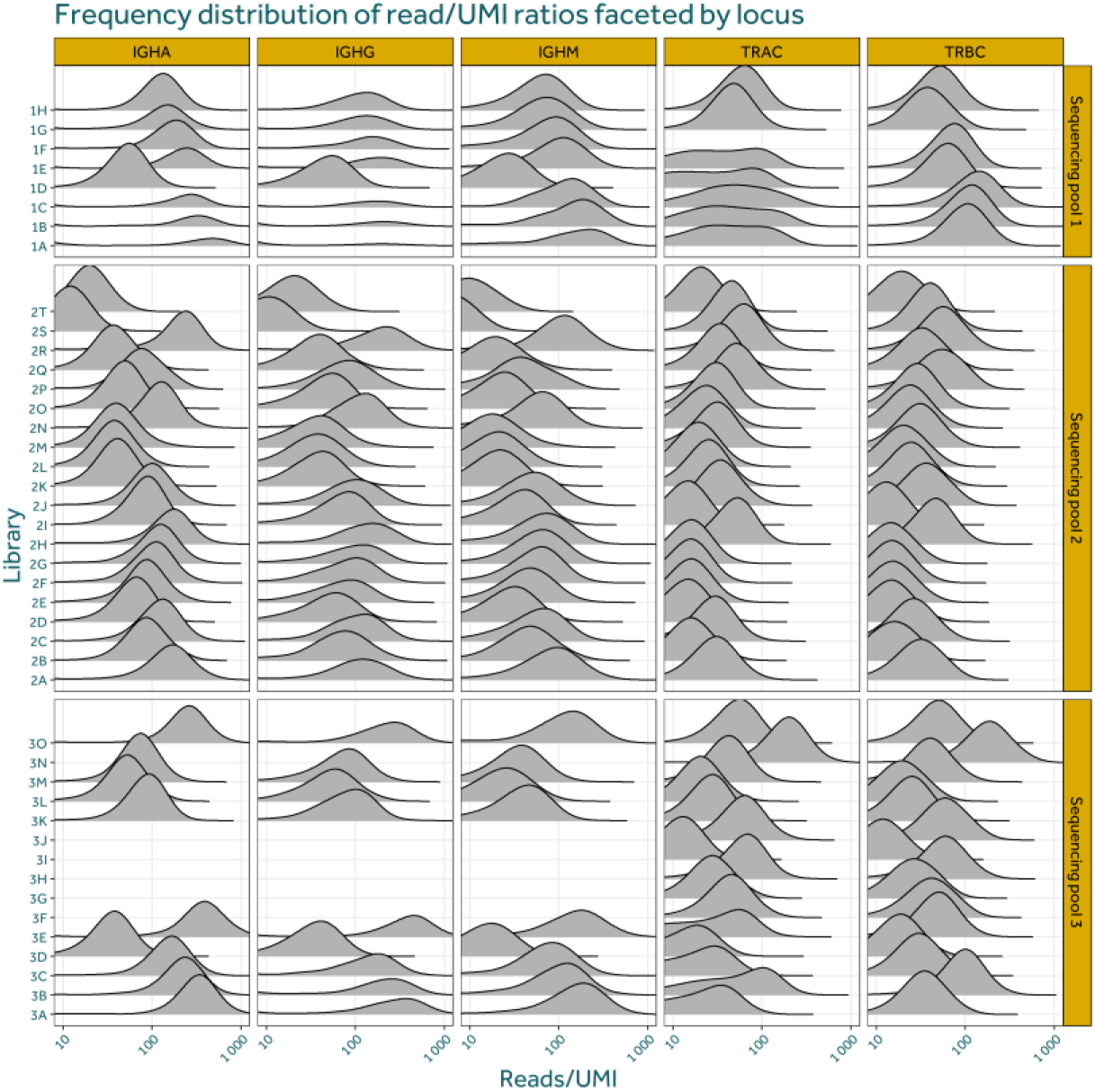
Reads per UMI distributions used to assess sequencing depth in DHIM1 libraries. Reads per UMI distributions derived from deeply sequenced, equimolarly pooled, and nf-core/airrflow assembled DHIM1 repertoires. Only sequences represented by a single UMI (duplicate_count = 1) were included, such that the reported consensus_count directly reffects the number of reads assigned to that UMI. These data indicate that sufficient reads per UMI coverage has been achieved to allow reliable enumeration of total UMI counts for subsequent Ct₂⁄₃–UMI calibration.

Figure 2 further shows that despite deep sequencing, IGH libraries 2T and 2S had IGHM read/UMI modes under or equal to ten, indeed demonstrating the intrinsic weakness of equimolar pooling of samples with varied size. As the UMI counts of these samples could be an underestimation and therefore unrepresentative, they were removed from further analysis to ensure accurate UMI-*Ct*_2/3_ modelling. For each sample, the total UMI-counts were determined by summing the duplicate count values of all sequences in the assembled repertoire. A log-linear regression of total UMI counts versus *Ct*_2/3_ values demonstrated an exponential relationship consistent with theoretical amplification kinetics (Figure 3A). The fitted model,

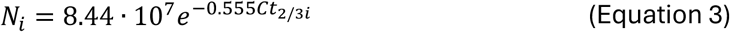

**Figure 3 –.**
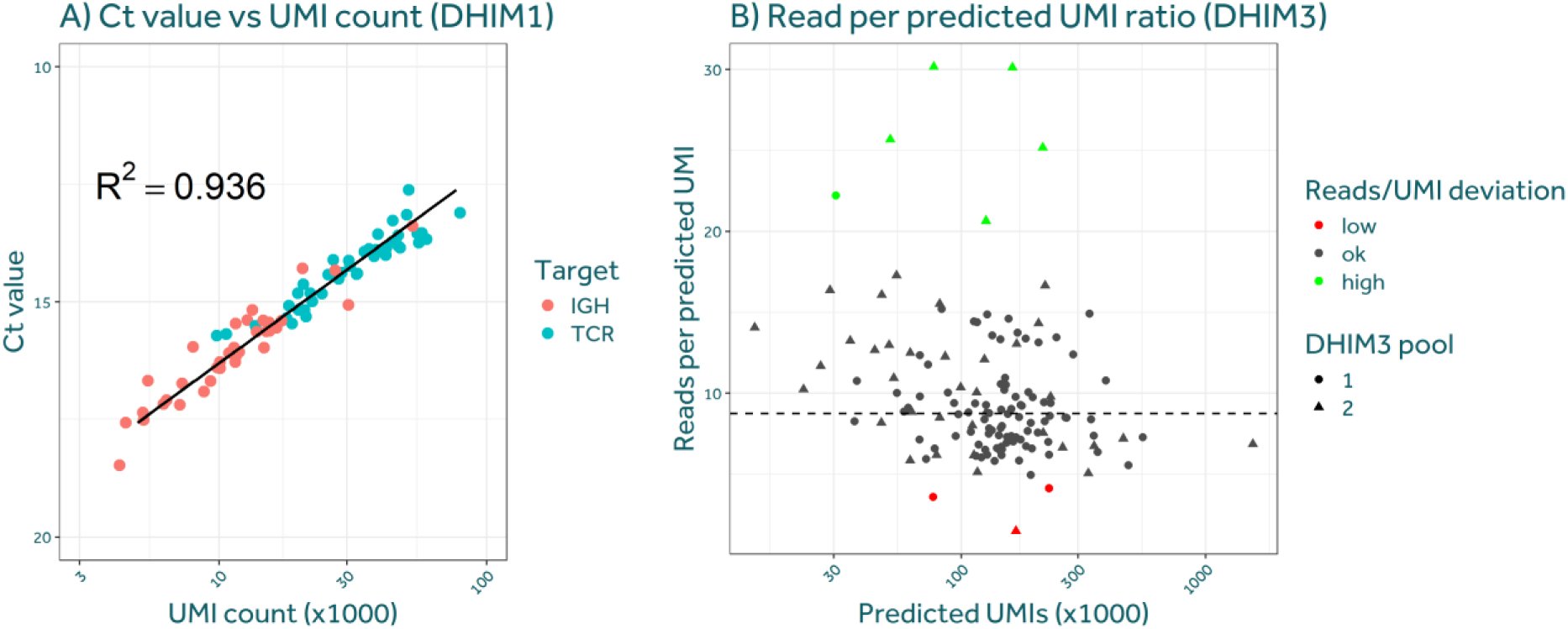
Ct–based UMI estimation and equi-depth pooling. (A) Relationship between PCR2 cycle threshold values (*Ct*_2/3_) and total UMI counts determined from deeply sequenced DHIM1 libraries. UMI counts were obtained by summing duplicate_count values from nf-core/airrffow–assembled repertoires after exclusion of under-sequenced libraries, ensuring reliable UMI enumeration. A log-linear regression was fitted to model the exponential relationship between amplification cycle number and detected UMI abundance. (B) Assessment of library pooling accuracy based on *Ct*_2/3_-derived UMI estimates in the independent DHIM3 cohort. Expected UMI counts obtained from the calibration model were compared with the number of sequencing reads assigned to each library after pooling, verifying that libraries were pooled proportionally to their estimated UMI abundance during the wet-lab pooling step. Point shapes indicate pool membership. Samples with reads per expected UMI at least twofold higher or lower than the cohort median (dashed line) are coloured green or red respectively.

showed a strong correlation (R² ≈ 0.94), validating *Ct*_2/3_ as a reliable proxy for library UMI abundance.

### Validation of the *Ct*_2/3_–UMI model to clinical samples from an independent study

The derived regression was then applied to an independent cohort from a DHIM3 study, which included longitudinal peripheral-blood samples from which B- and T-cell repertoire libraries were prepared. For each library, the *Ct*_2/3_ value was used to predict relative UMI abundances, and the samples were distributed across two equi-depth pools (STAR★METHODS).

Following sequencing, the number of reads assigned to each sample was evaluated (Figure 3B). Overall, the equi-depth pooling strategy resulted in a largely uniform read per expected UMI ratio across libraries, indicating that sequencing effort was successfully normalized to expected sample complexity. Minor but appreciable deviations from the target ratio were observed, consistent with technical variability during the pooling step, most plausibly due to pipetting inaccuracies. A small subset of samples received read allocations that were at least twofold higher or twofold lower than the cohort median and are highlighted in green (oversequenced) or red (undersequenced).

To assess the impact of sequencing depth on immune receptor UMI recovery, we first examined the read support of immune receptor sequences represented by a single UMI in the assembled DHIM3 repertoires (Figure 4A). For these sequences (duplicate_count = 1), the consensus_count corresponds to the number of reads supporting that individual UMI-derived consensus sequence. Figure 4A shows, for each sample and DHIM3 pool, the proportion of single- UMI immune receptor sequences observed at each read/UMI value, after excluding single-read consensus sequences. The distributions were summarized across all retained immune receptor categories and separately by isotype/receptor class, with curves coloured according to whether the sample-level reads/UMI value was below, within, or above the expected range. Overall, most single-UMI immune receptor sequences were supported by low read counts, with higher read/UMI values enriched in samples classified as oversequenced.

**Figure 4 –.**
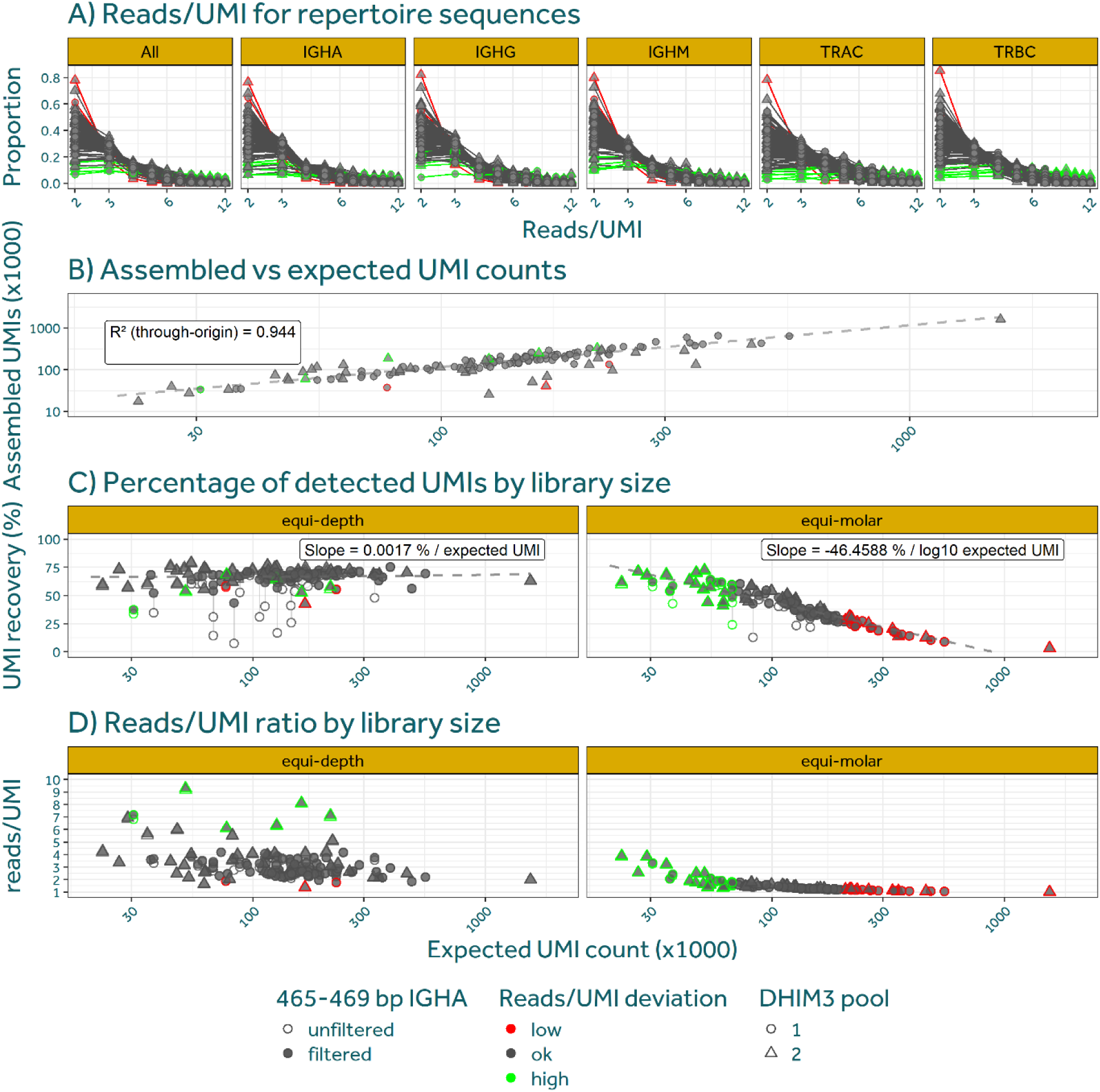
Impact of equi-depth pooling on effective sequencing depth and UMI recovery in the DHIM3 cohort. The same sample-specific point shapes are used consistently across all panels and indicate pool membership (Pool 1, Pool 2). Samples with a read/expected UMI ratio at least twofold higher or lower than the cohort median are coloured green or red respectively. (A) Read-per-UMI distributions for target-specific immune receptor sequences represented by a single UMI (duplicate_count = 1) in assembled repertoires, stratified by locus. (B) Assembled UMI counts plotted against *Ct*_2/3_-derived expected UMI counts. Assembled UMIs scale approximately linearly with expectations, indicating that the Ct-based model captures relative library sizes. Systematic deviations reffect differences in achieved read-per-UMI ratios, with higher ratios yielding slightly increased UMI recovery and lower ratios yielding reduced recovery, underscoring the importance of maintaining comparable read/UMI ratios across samples. (C-D) UMI recovery and reads per UMI under equi-depth pooling versus an equimolar-like subsampling comparator. Panels are faceted by pooling strategy. In each panel, unfiltered data are shown as open points and data filtered for the 4c5–4cS bp IGHA off-target band as filled grey points. Grey dashed lines connect repeated measurements of the same sample. (C) UMI recovery, defined as the fraction of observed UMIs relative to Chao1-estimated richness, plotted against expected UMI counts. Dashed regression lines are fitted to the filtered data per strategy: ordinary least squares on expected UMI for equi- depth, and on log10-transformed expected UMI for equimolar. Slope estimates are shown as labels. The near-ffat regression under equi-depth pooling indicates stable UMI recovery across library sizes. The negative log-linear slope under equimolar subsampling indicates that larger libraries are systematically undersampled when reads are allocated uniformly. (D) Mean reads per UMI as a function of expected UMI count. Equi-depth pooling maintains broadly constant sequencing depth per molecule across small and large libraries once non-informative off-target products are excluded. Equimolar subsampling shows a marked decline in reads per UMI with increasing library complexity.

The predictive potential of the *Ct*_2/3_ values for immune receptor sequence recovery at these cost-effective read/UMI ratios was evaluated by plotting the assembled UMI counts against the expected UMI counts (Figure 4B). Assembled UMI counts increase approximately linearly with the expected UMI counts, with most libraries clustering close to the regression line through the origin, indicating that the Ct-derived model captures relative library sizes well.

To further dissect the sources of variation in effective sequencing depth, reads per UMI distributions were stratified by sequence length using the collapsed UMI tables generated immediately after read pairing and consensus construction (Supplementary Figure 1). In this analysis, sequence lengths were grouped into 5 bp bins. Within each sample, pool, and immune receptor/isotype category, each bin was summarized by its share of sequences and by the mean consensus read count per sequence, calculated separately before and after assembly/quality filtering. Colouring by mean consensus count enabled direct comparison of sequencing depth across length classes and samples.

Across libraries, the overall length abundance and reads per UMI profiles were consistent, indicating that the equi-depth pooling strategy resulted in comparable sequencing effort per molecule independent of the sample. However, clear differences were observed between length distributions of sequences retained in the assembled repertoires and those that were excluded. In the unassembled sequences, a sharp peak is observed in sequences amplified by the IGHA primer with a size of circa 465 bp. For this peak, the read/UMI ratio was around 1. As this peak is not observed in the assembled sequences, and is more abundant in libraries with few BCR or TCR UMI counts, it is consistent with non-specific amplicons that are co-amplified during PCR1 or PCR2 despite bead-based size selection. Because these products do not contribute to the final assembled repertoires, they are not relevant for evaluating effective sequencing depth or UMI recovery and must be excluded to avoid distorting downstream metrics. Therefore, IGHA- sequences of 465-469 bp were removed from the dataset for further analysis.

To test whether variation in library size was compensated by assigning more reads to larger libraries, and to benchmark the equi-depth strategy against conventional equimolar pooling, we quantified UMI recovery as a function of the expected UMI count predicted by the Ct-based model. UMI recovery was computed from the collapsed UMI tables. For each sample and primer- consensus group, we estimated the total UMI richness using Chao1 and expressed UMI recovery as the fraction of observed UMIs relative to this estimate. This operation was performed on both the equi-depth sequencing data and an equimolar-like subsampled comparator. Analyses were run on the unfiltered data and on the data from which the non-specific IGHA 465–469 bp off-target peak was removed (Figure 4C, open and filled points respectively). For the filtered data, ordinary least squares regression was fitted separately per pooling strategy. Under equi-depth pooling, UMI recovery remained flat across the range of library sizes (slope = 0.0017 % / 1000 expected UMI, RMSE = 6.38%), whereas the equimolar subsampled data exhibited a negative slope with respect to log10-transformed expected UMI count, indicating that larger libraries were systematically undersampled under uniform read allocation.

In parallel, sequencing depth per molecule was evaluated by plotting the mean reads per UMI against the expected UMI count for each sample (Figure 4D ), again using collapsed UMI tables because they preserve singleton UMIs and therefore provide an unbiased view of shallow coverage. Under equi-depth pooling, mean reads per UMI remained broadly constant across library sizes. In contrast, the equimolar subsampled comparator showed a marked decline in reads per UMI with increasing expected UMI count, reflecting the dilution of sequencing effort across the most complex libraries under uniform read allocation. Taken together, Figure 4C and Figure 4D demonstrate that differences in library size, quantified by expected UMIs, are effectively compensated by equi-depth read allocation: across small and large libraries, both the Chao1- based percentage of detected UMIs and the mean read-per-UMI depth remain broadly comparable once non-informative off-target products are excluded, whereas equimolar-like allocation produces a complexity-dependent bias. However, the difference between the unfiltered and filtered analyses also shows that Ct-based UMI estimates can be biased when non-specific amplicons contribute substantially to the qPCR signal or collapsed UMI counts. Therefore, for new sample types, primer sets, or library-preparation chemistries, a pilot sequencing experiment may be required to identify target-specific amplicons and establish an appropriate Ct-to-UMI calibration, as discussed below.

### Cost saving estimations

The central hypothesis of this study is that equimolar pooling is intrinsically inefficient for heterogeneous AIRRseq sample pools, because it allocates sequencing reads according to library molarity rather than molecular complexity, thereby forcing the total sequencing depth to be determined by the largest library in the pool. The cost-saving analyses presented here test this hypothesis by quantifying between-sample dispersion in UMI abundance and translating this dispersion into expected relative sequencing requirements.

The relative reduction in sequencing effort achieved by equi-depth pooling is determined by:

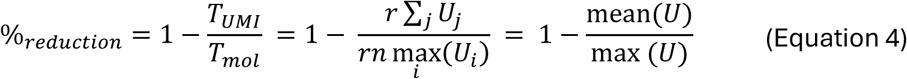

We next quantified the distribution of library sizes in the DHIM3 cohort. The per-sample UMI-count distribution is shown in Figure 5A (n = 136 libraries; mean 188k UMIs; max 1,634k UMIs; 94-fold range). Applying (Equation 4 to the DHIM3 cohort shows that equimolar pooling would require approximately 2 billion reads to achieve 9 reads per UMI across all libraries, compared to 230 million reads under equi-depth pooling; a reduction of 89%. This discrepancy is driven by one sample with an exceptionally high UMI count (∼9-fold above the cohort mean), which under equimolar pooling sets the sequencing depth for all other samples. In practice, such an outlier could be mitigated under equimolar pooling by assigning it to a separate pool or by selective re- sequencing. Nevertheless, the observed reduction is indicative of the savings potential, because the DHIM3 cohort was generated under controlled experimental conditions. Real-world clinical batches typically exhibit greater variance in library sizes, which would further increase the cost differential.

**Figure 5 –.**
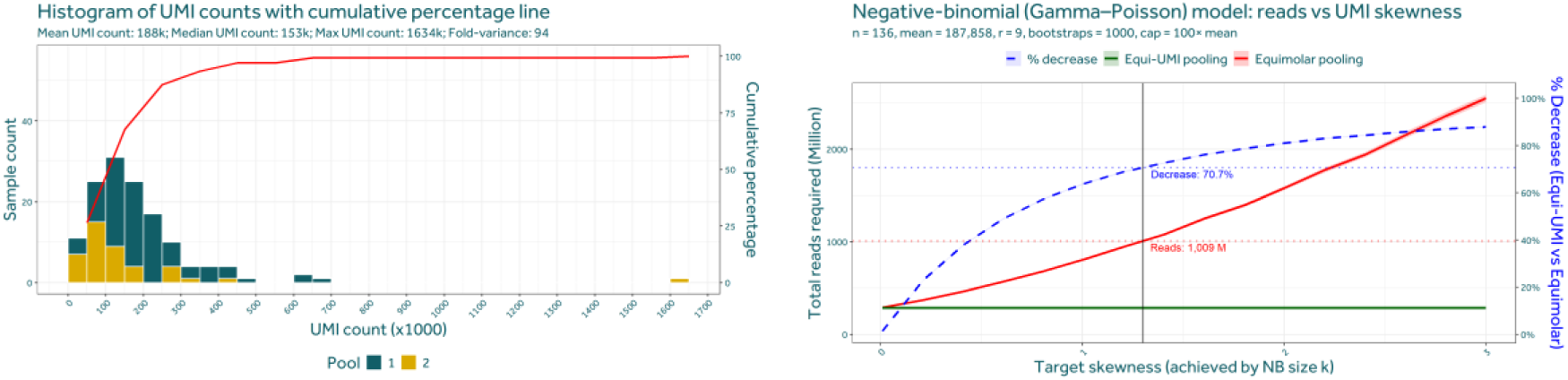
Quantifying and generalizing sequencing cost savings achieved by equi-depth pooling. (A) Distribution of per-sample UMI counts in the DHIM3 cohort. Bars show the number of samples per UMI-count bin, and the red line indicates the cumulative percentage of samples. The distribution spans a S4-fold range between the smallest and largest libraries and exhibits a right-tailed, negative-binomial–like shape, indicating substantial between-sample dispersion in molecular complexity. (B) Model-based estimation of relative sequencing requirements for equimolar versus equi-depth pooling as a function of UMI-count distribution skewness. For each target skewness, UMI-count distributions were simulated using a negative-binomial (Gamma–Poisson) model, and the total reads required to achieve a S reads-per-UMI depth were computed for both pooling strategies. Solid lines indicate mean requirements across bootstrap replicates, with shaded ribbons denoting S5% confidence intervals. The dashed blue line (right axis) shows the corresponding percentage reduction in required reads achieved by equi-depth pooling relative to equimolar pooling. The vertical line marks the skewness inferred from the DHIM3 cohort, illustrating that the observed sample heterogeneity lies in a regime where equi-depth pooling yields substantial (∼70%) and predictable sequencing cost reductions.

To generalize these findings beyond a single cohort, it is necessary to explicitly model the distribution of library sizes within a sequencing pool. As a first step, we examined how variance in information content depends on biological target. Supplementary Figure 2A shows UMI-count distributions for DHIM1 samples stratified by target locus. These data demonstrate that dispersion is strongly target-dependent and reflects underlying biological processes. In particular, IGH libraries exhibit pronounced right-tailed distributions with occasional extreme UMI counts, consistent with massive plasmablast expansions that are characteristic of acute dengue infection. In contrast, TCR libraries show a more flat distribution, indicating more limited between-sample variability.

The distribution of library sizes in the DHIM3 cohort (Figure 5A) follows a negative-binomial (Gamma–Poisson)–like shape, with most samples clustered near the mean and a minority of samples contributing a disproportionate share of total UMIs.

Supplementary Figure 2B illustrates the shape of negative-binomial UMI distributions across a range of skewness values, with the mean normalized to one. Increasing skewness produces progressively heavier right tails and increases the separation between typical and extreme library sizes. Using a skewness value of approximately 1.3, we generated simulated sample-size distributions that closely resemble the empirical DHIM3 distribution, as shown in Supplementary Figure 2C. This indicates that the DHIM3 cohort is representative of a moderately skewed but biologically realistic sample pool.

Finally, using these simulation parameters, we estimated the expected relative cost savings of equi-depth pooling as a function of distribution skewness. Figure 5B summarizes these results and shows that the advantage of equi-depth pooling increases monotonically with skewness, reflecting the growing divergence between the mean and the maximum library size. The empirically observed 89% reduction in the DHIM3 cohort exceeds the simulated estimate at the cohort’s skewness (approximately 70%), because the empirical maximum UMI count (1,634k) was ∼9-fold above the mean, placing it beyond the range captured by the fitted negative-binomial model. Together, these analyses demonstrate that substantial sequencing cost reductions are an expected and predictable consequence of equi-depth pooling in heterogeneous AIRRseq sample pools.

## Discussion

This study demonstrates that equi-depth pooling based on *Ct*_2/3_-derived UMI estimates provides a practical and robust solution to one of the main inefficiencies in bulk AIRRseq experiments: the mismatch between sequencing effort and underlying molecular complexity. By allocating sequencing reads in proportion to expected UMI abundance rather than library molarity, equi-depth pooling aligns sequencing depth with the number of informative molecules per sample and thereby improves both cost efficiency and quantitative consistency across heterogeneous clinical libraries.

### Target-specific and unspecific UMI-counts

5’ RACE library preparation relies on reverse primers for target specificity. For BCR and TCR amplification, these primers are partially degenerate to enable amplification across diverse alleles. As a consequence, substantial amounts of non-target amplicons are generated, even after bead-based size selection. The composition and relative abundance of these off-target sequences vary between library preparation kits and sample types, and they are amplified with efficiencies that differ from those of target-specific amplicons. This results in systematically different reads per UMI ratios for target and non-target sequences (Supplementary Figure 1). Therefore, a single reads per UMI value cannot be meaningfully assigned to an entire sequencing library. Instead, target-specific sequences must first be identified, after which their reads per UMI ratios can be estimated.

Using the nf-core/airrflow pipeline, two strategies are available to estimate sequencing depth and UMI coverage while restricting the analysis to target-relevant sequences.

The first strategy is to perform reads per UMI estimation on the final assembled repertoires. In these outputs, the pipeline reports the number of reads per consensus sequence in the consensus_count column and the number of UMIs per sequence in the duplicate_count column. For sequences with a duplicate count of one, the consensus count directly reflects the number of reads assigned to a single UMI. The distribution of these values therefore provides an estimate of UMI coverage for target-specific sequences (Figure 2 and Figure 4A), while the total UMI count per sample can be approximated by summing duplicate_count values. However, these repertoires have undergone extensive quality control and filtering, meaning they are no longer representative of the raw sequencing data. In particular, sequences supported by only a single read are removed by the pipeline. In cases of under-sequencing, or when only a fraction of UMIs is captured, this filtering prevents accurate estimation of the true UMI count, limiting the applicability of diversity estimators such as Chao1.

The second strategy uses intermediate outputs from the assembly process, specifically the collapsed UMI tables generated after read pairing and consensus building. These tables report the read count per UMI and the length of the assembled sequence (Supplementary Figure 1), and they retain UMIs supported by a single read. In this approach, sequences are filtered based on the presence of target-specific primer sequences, ensuring enrichment for BCR and TCR amplicons while preserving singleton UMIs. This enables more accurate estimation of total UMI counts and reads per UMI ratios, particularly for shallowly sequenced libraries. The main limitation of this strategy is that target specificity is inferred solely from primer presence. No additional confirmation, for example via BLAST-based annotation, is performed at this stage, and residual non-specific sequences may therefore remain and bias the estimated read per informative UMI values.

Together, these two approaches represent a trade-off between specificity and representativeness. Analyses based on assembled repertoires provide high-confidence target sequences but underestimate UMI diversity in under-sequenced libraries, whereas analyses based on collapsed UMI data preserve singleton information at the cost of potentially including non-target amplicons.

### Pipetting likely contributes to sequencing depth variance

Figure 3B shows that the achieved reads-to-expected-UMI ratios can differ appreciably between libraries. Likely, this variability reflects technical variation introduced during the wet-lab pooling step, most plausibly due to pipetting inaccuracies when combining microliter-sized library volumes. As a result, this variability represents technical noise of the pooling procedure rather than a limitation of the Ct-based UMI prediction model. The variability is consistent across both Pool 1 and Pool 2.

Crucially, this pooling-induced variability explains most of the downstream variation observed across multiple analyses. The spread in reads per UMI for assembled immune receptor sequences in Figure 4A closely mirrors the variation seen in reads per expected UMI in Figure 3B, indicating that differences in effective sequencing depth are primarily inherited from the pooling step and that the true per-sample UMI diversity correlates with the expected diversity. The same pattern is observed in Figure 4B, where deviations between assembled and expected UMI counts align with over- or under-sequencing relative to the target read/UMI ratio. Similarly, variation in the fraction of detected UMIs in Figure 4C is largely attributable to this technical variability, rather than incorrect underlying UMI diversity estimations. Together, these observations demonstrate that once true biological differences in UMI abundance are accounted for by equi-depth pooling, residual variation in sequencing outcomes is dominated by pipetting error during pooling.

### Equi-depth pooling mitigates sampling biases

UMI recovery is remarkably stable across the full range of library sizes when equi-depth pooling is applied. The near-flat relationship between Chao1-based UMI recovery and expected UMI count in Figure 4C under equi-depth pooling indicates that allocating proportionally more reads to larger libraries successfully compensates for differences in molecular complexity. This flat slope is the expected outcome if sequencing effort is correctly normalized to library size and is evidence that the pooling strategy mitigates sampling biases. In direct contrast, the equimolar subsampled comparator shows a pronounced negative relationship between UMI recovery and library size, confirming that conventional uniform read allocation systematically undersamples the most complex libraries. This comparison quantifies the sampling bias that equi-depth pooling is designed to prevent.

### Over-sequencing increases non-specific reads

The IGHA off-target amplicons observed in Supplementary Figure 1 are characterized by very low read/UMI ratios and therefore inflate singleton counts, biasing richness estimates in unfiltered data. Once these non-informative products are removed, UMI recovery becomes largely independent of library size (Figure 4C). Notably, samples with higher read-to-expected-UMI ratios (Figure 3B) showed lower estimated recovery in the unfiltered analysis, an effect that was largely resolved by filtering. This observation suggests that excess sequencing depth can preferentially increase detection of non-specific products rather than improve recovery of biologically relevant immune receptor UMIs, reinforcing that oversampling, which will disproportionately affect low diversity samples in equimolar pooling, is not only inefficient but can actively distort downstream diversity estimates.

### Relative UMI-estimations are replicable across protocol differences

A consistent observation was that assembled UMI counts in the DHIM3 cohort exceeded the Ct-based predictions by approximately 50 percent (Figure 4B). Importantly, this deviation was uniform across samples, indicating a systematic offset rather than sample-specific technical noise. Such offsets are expected when calibration and validation datasets differ in library-preparation chemistry, RNA input amount, or primer composition, all of which can introduce a global scaling factor in the relationship between Ct values and absolute UMI counts.

Crucially, the Ct-based model is intended to capture relative differences in UMI diversity between samples, not to provide universally transferable absolute UMI estimates. In this respect, the model performed as designed: assembled UMI counts scaled linearly with expected UMI counts across the DHIM3 libraries, demonstrating that relative library sizes were accurately recovered independent of target locus or sample complexity. This confirms that Ct-derived estimates robustly preserve the ordering and proportional differences in UMI abundance between samples.

## Conclusions

Taken together, these results show that equi-depth pooling, as compared to equimolar pooling, improves the interpretability and comparability of AIRRseq data by stabilizing read/UMI ratios across samples and can potentially deliver substantial cost savings. By preventing systematic under-sequencing of large libraries and oversampling of small ones, this approach reduces technical bias in UMI recovery, diversity estimation, and downstream quantitative analyses. In cohorts characterized by extreme biological heterogeneity, such as acute viral infections with transient plasmablast expansions, these advantages are particularly pronounced. More broadly, equi-depth pooling provides a cost-effective, simple, scalable strategy that can be readily integrated into existing RNA-based UMI workflows and adapted to other library-preparation chemistries where sequencing depth must be matched to molecular complexity.

## Resource availability

### Lead contact

Requests for further information and resources should be directed to and will be fulfilled by the lead contact, Thomas-Wolf Verdonckt.

### Materials availability

This study did not generate new unique reagents.

### Data and code availability

All data necessary to replicate the published results will be provided as supplementary files, and all original code will be published on GitHub, upon acceptance in a peer-reviewed journal.

## Acknowledgements

Funding: this work was supported by grants from: Agency of Innovation and Entrepreneurship of the Flemish Government (HBC.2022.0988) and Johnson and Johnson;

## Author contributions

Conceptualization: TWV, MS, KB, KKA; Methodology: TWV, MS, KB; Investigation: TWV, MS; Visualization: TWV; Funding acquisition: FVN, OL, KKA; Project administration: FVN, OL, KKA; Supervision: FVN, OL, KKA; Writing – original draft: TWV; Writing – review & editing: TWV, KB, ASV, CS, MS, FVN, OL, KKA;

## Declaration of interests

Authors declare that they have no competing interests.

## Declaration of generative AI and AI-assisted technologies

During the preparation of this work the author(s) used ChatGPT to screen the literature and to improve language and readability. After using this tool/service, the author(s) reviewed and edited the content as needed and take(s) full responsibility for the content of the published article.

## STAR★METHODS

### KEY RESOURCES TABLE

| Reagent or resource | Source | Identifier |
| --- | --- | --- |
| PAXgene® RNA tubes (IVD) | QIAGEN | cat. no. 762165 |
| PAXgene® Blood RNA Kit (IVD) | QIAGEN | cat. no. 762164 |
| Qubit™ RNA HS Assay Kit | Invitrogen | Q32852 |
| Qubit 4 Fluorometer | Invitrogen | Q33238 |
| TapeStation RNA ScreenTape & Reagents | Agilent | 5067 |
| 4200 TapeStation System | Agilent | G2991BA |
| Zymo RNA Clean & Concentrator™-5 | Zymo Research | R1013 |
| NEBNext® Immune Sequencing Kit (Human) | New England Biolabs | E6320S |
| CFX96™ Real-Time System | Bio-Rad | 1845097 |
| KAPA SYBR FAST Master Mix Universal (2x) | Roche | KK4601 |
| NEBNext Library Quant DNA Standards | New England Biolabs | E7642S |
| Tween-20 | Merck | P1379 |

### PRIMERS TABLE

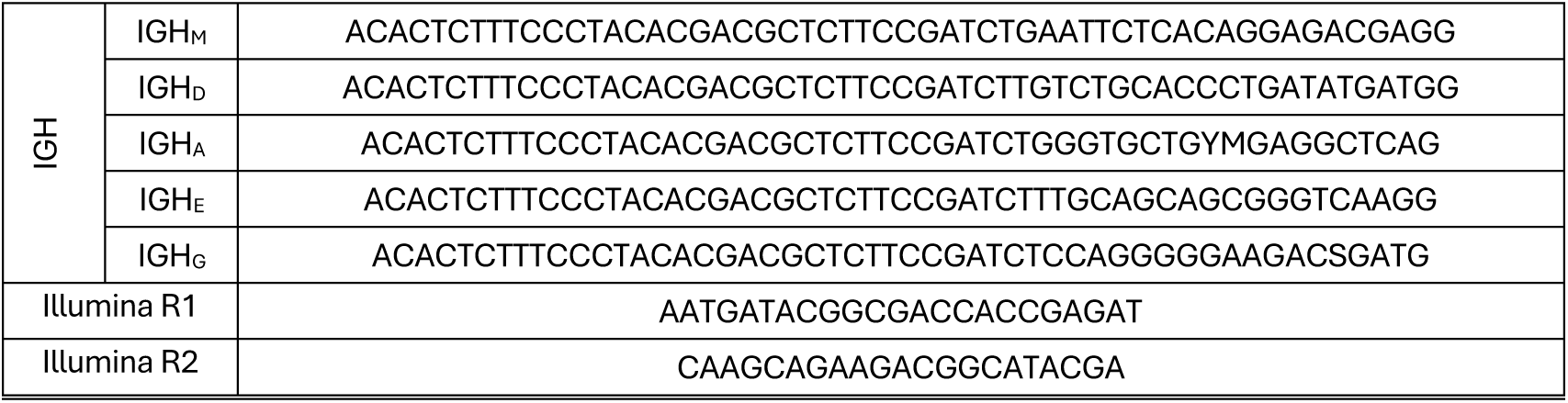

### EXPERIMENTAL MODEL AND STUDY PARTICIPANT DETAILS

This study involved the secondary analysis of biobanked total RNA obtained during a Dengue-1- Virus Live Virus Human Challenge study (DENV-1-LVHC; ClinicalTrials.gov ID NCT03869060)^15^, together with whole-blood samples collected in a trial evaluating the prophylactic activity of JNJ- 64281802 during a Dengue-3-Virus Live Virus Human Challenge (DENV-3-LVHC; ClinicalTrials.gov ID NCT05048875).

The DENV-1-LVHC protocol received approval from the State University of New York Upstate Medical University Institutional Review Board (SUNY-UMU; 1252992) and the Department of Defense’s Human Research Protection Office. The DENV-3-LVHC protocol was approved by independent ethics committees at the Johns Hopkins Bloomberg School of Public Health and the University of Vermont. All participants in both studies provided written informed consent before any samples or data were collected.

The use of samples and associated personal or medical data from the DENV-1-LVHC and DENV- 3-LVHC studies for the present analysis was reviewed and approved by the Institutional Review Board of the Institute of Tropical Medicine Antwerp under reference 1776/24 and by the Ethics Committee UZA/UAntwerp under reference 6738.

An overview of participant demographics and relevant characteristics is provided in Table 1.

**Table 1 –.**
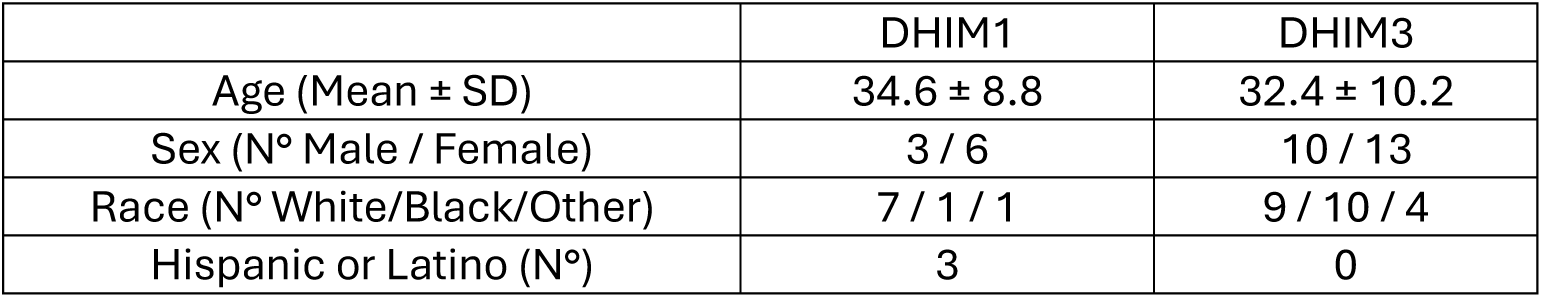
Participant demographics and characteristics of the dengue human infection model studies.

### METHOD DETAILS

This protocol describes how to generate immune repertoire sequencing libraries with the NEBNext Immune Sequencing Kit (Human) following a modified version of the manufacturer’s manual^14^ (version 2.0_12/21) and pool them so that, after sequencing, libraries achieve a comparable read depth relative to library complexity, operationalized as a similar read-per-UMI ratio across samples. The method uses information already generated during standard library preparation, specifically the cycle number corresponding to two-thirds of maximal fluorescence (*Ct*_2/3_) obtained during the qPCR step (step 5.4 of the instruction manual), combined with library molarity measurements, to determine pooling volumes.

Although the exact implementation described here is specific to the NEBNext Immune Sequencing Kit (Human), the working principle can be applied to other library preparation chemistries that include a PCR amplification step to a target yield, provided that a calibration model linking *Ct*_2/3_ (or an equivalent amplification metric) to library complexity has been established as described in the manuscript.

#### Total RNA extraction from PAXgene® Blood RNA tubes and quality control

Whole-blood samples (2.5 ml) were collected in PAXgene RNA tubes (QIAGEN) and the RNA extracted through the PAXgene® Blood RNA Kit (QIAGEN) according to the PAXgene® Blood RNA Kit Instructions for Use (Handbook Version 2, February 2023). The extracted RNA was quantified on a Qubit™ High Sensitivity RNA Assay (Thermo Fisher Scientific) and size profiles were obtained through RNA ScreenTapes on the TapeStation system (Agilent). When necessary, the RNA samples were concentrated to ≥110 ng/µl with the Zymo RNA Clean C Concentrator™-5 kit (Zymo Research).

#### Sequencing library preparation with the NEBNext Immune Sequencing Kit (Human)

This protocol includes mandatory modifications to the manufacturer’s NEBNext Immune Sequencing Kit (Human) workflow^16^. Unless explicitly stated below, all steps follow the NEBNext instruction manual (Version 2.0, 12/21). These modifications must be applied exactly as described. For each sample, 1 µg of total RNA was used for cDNA synthesis. The resulting cDNA was split into three volumes, and bulk TCR and IGH libraries were prepared in parallel using one volume each.

##### Modification 1: Additional denaturation step prior to cDNA synthesis

In contrast to the manufacturer protocol, reverse transcription reagents are not added in a single master mix. Instead, RNA is first combined with primers and dNTPs and subjected to a denaturation and rapid quenching step before addition of the remaining reverse transcription components. Replace step 1.1 of the NEBNext instruction manual (Version 2.0, 12/21) with the following steps:

1. Mix the following components in a sterile nuclease-free tube:

| Component | Volume (µl) |
| --- | --- |
| Total RNA | 1-9 |
| ● (lilac) NEBNext IS RT Primer | 1 |
| ● (lilac) dNTP Solution Mix | 2 |
| ○ (white) Nuclease-free Water | Variable |

2. Set a 100 µl or 20 µl pipette to 12 µl and then pipette the entire volume up and down at least 10 times to mix thoroughly. Perform a quick spin to collect all liquid from the sides of the tube.
3. Place in a thermal cycler, with the heated lid set to ≥80°C, and incubate for 5 minutes at 65°C.
4. Immediately transfer the tubes to ice-cold water and actively move the tubes within the water to ensure rapid and homogeneous cooling.
5. Proceed immediately to cDNA synthesis by adding the remaining reverse transcription components:

| Component | Volume (µl) |
| --- | --- |
| ● (lilac) NEBNext IS RT Buffer (4x) | 5 |
| ● (lilac) NEBNext IS TS Oligos | 1 |
| ● (lilac) NEBNext First Strand Synthesis Enzyme Mix | 2 |
| ● (lilac) NEBNext Cell Lysis Buffer | 0.5 |
| Total Volume | 20.5 |

6. Proceed with step 1.2 of the NEBNext instruction manual

Critical note:

Failure to perform this denaturation and rapid quenching step can result in inefficient cDNA synthesis or dominance of non-specific amplification products. This leads to low BCR- and TCR-sequence recovery, and unreliable *Ct*_2/3_ values during PCR2 qPCR compromising downstream equi-depth pooling.

##### Modification 2: IGH primer mix

In step 3.1 of the manufacturer protocol, the NEBNext IS BCR Primers (Human)(NEB # E2626AA), are replaced by a primer mix that targets only the heavy-chain sequences (IGH, primer table). Each primer is mixed to a concentration of 1 µM.

###### Critical note

The immunoglobulin heavy chain (IGH) and light chain (IGLκ and IGLλ) constant-region primers used in the NEBNext IS BCR Primers (Human)(NEB # E2626AA) exhibit different amplification efficiencies. The BCR primer mix therefore produces amplification products with unequal copy numbers. As a result, the *Ct*_2/3_ value cannot be interpreted as a reliable proxy for the complexity of both heavy and light chain libraries simultaneously, and it is not possible to obtain a homogeneous read-per-UMI ratio for all target types within and between samples. If light-chain sequences are desired, a separate library preparation should be carried out using only light-chain primers.

##### Modification 3: *Ct*_2/3_ estimation

For accurate assessment of UMI counts, the *Ct*_2/3_ value determination should be performed computationally in replicate. The qPCR step for PCR2 cycle optimization was performed on a CFX96™ Real-Time System. 10 µl of the mixture obtained in step 5.2 of the manufacturer protocol was pipetted into two wells of a 96-well plate for increased accuracy. To determine the *Ct*_2/3_ values, the raw data was exported in .csv format and processed using a custom script.

#### Sequencing library quantification

The obtained libraries were quantified through qPCR using NEBNext Library Quant DNA Standards (New England Biolabs). First, 2 µl of the libraries were diluted to the 10 nM range by mixing 1:2000 in 0.1% Tween-20 through serial dilution. Next, the samples and standards were run in triplicate on a CFX96™ Real-Time System. For each well, the following mixture was used:

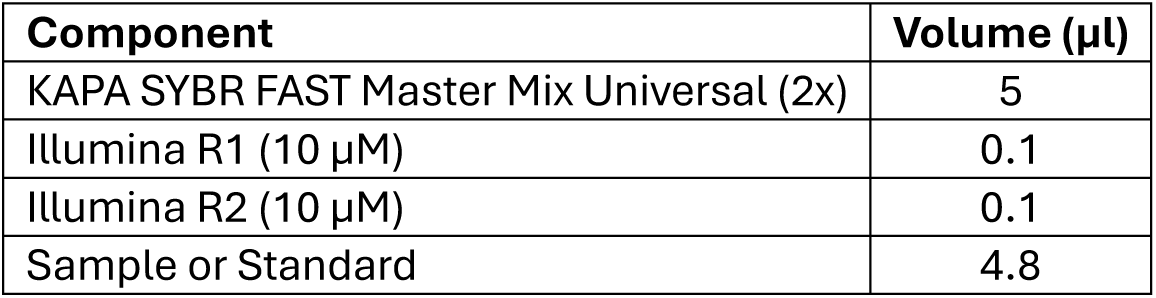

| Component | Volume (µl) |
| --- | --- |
| KAPA SYBR FAST Master Mix Universal (2x) | 5 |
| Illumina R1 (10 µM) | 0.1 |
| Illumina R2 (10 µM) | 0.1 |
| Sample or Standard | 4.8 |

The following PCR cycle was used:

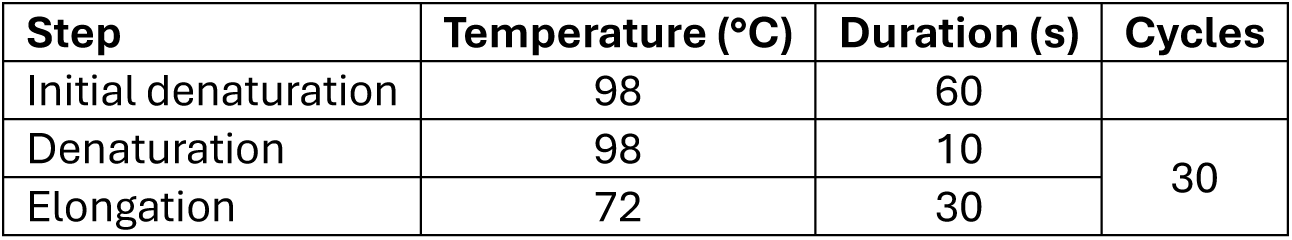

The raw data was exported in .csv format. The standard curve regression, quality control and absolute sample quantification were performed using a custom script, accounting for amplicon size.

#### Equi-depth pooling

The *Ct*_2/3_ values were used to compute the expected UMI count of each sample through the fitted model (Equation 3). Next, to obtain an equi-depth sequencing pool, the contribution of each sample to the pool can be determined through:

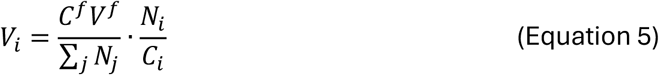

where *N_i_* is the UMI count of sample *i*, *C_i_* is the measured library concentration of sample *i*, ∑*_j_ N_j_* is the total UMI count across all samples, *C^f^* is the desired final pool concentration, and *V^f^* is the desired final pool volume.

In the DHIM3 experiment, samples were distributed across two independent equi-depth pools (Pool 1 and Pool 2) to accommodate the number of libraries and sequencing depth. The pooling formula was applied independently within each pool.

For pools with moderate variance in sample concentrations and expected UMI counts, the required sample volumes will be within a pipettable volume range (lower threshold of ∼1 µl determined by pipet accuracy, and upper threshold of ∼15 µl determined by library volumes) and the samples can be pooled ‘directly’, adding nuclease-free water to bring the pool to the final volume *V^f^*. However, for pools with large sample variance, the required volumes might differ too greatly requiring intermediate pooling steps. The expected UMI counts, pooling strategy, and required sample volumes were determined with a custom script.

The resulting pool was circularized and sequenced on an AVITI M flowcell (Element Biosciences) as previously described^16^.

#### Library assembly

Sequenced reads were processed with the nf-core/airrflow pipeline (version 5.0.0) on the CalcUA cluster (VSC, UAntwerp) using Nextflow and Apptainer for containerization. Separate runs were executed for IGH and TCR libraries using the nebnext_umi_bcr and nebnext_umi_tcr profiles respectively, which configure the pipeline for NEBNext Immune Sequencing library formats with UMI-based deduplication. Clonal analysis was disabled for all runs. Raw paired-end reads were quality-trimmed, UMI-corrected, and assembled into consensus sequences using the Immcantation framework. The pipeline outputs collapsed duplicate tables (retaining singleton UMIs) and final assembled repertoire files in AIRR Community standard format, which were used for downstream UMI counting, Chao1 richness estimation, and read-per-UMI calculations.

#### Equimolar-like subsampling comparator

To provide a direct experimental comparison between equi-depth and equimolar pooling strategies, an equimolar-like subsampling analysis was performed on the DHIM3 libraries. For each library, sequencing reads were randomly subsampled to 200,000 reads per sample, mimicking the uniform read allocation that results from equimolar pooling. Subsampled libraries were re-processed through the nf-core/airrflow analysis pipeline version 5.0.0 to generate independent assembly outputs. UMI recovery (Chao1-based) and reads-per-UMI metrics were then computed from the subsampled collapsed UMI tables using the same procedures applied to the equi-depth data, enabling direct comparison in Figure 4C and Figure 4D.

## Supplementary data

**Supplementary Figure 1 –.**
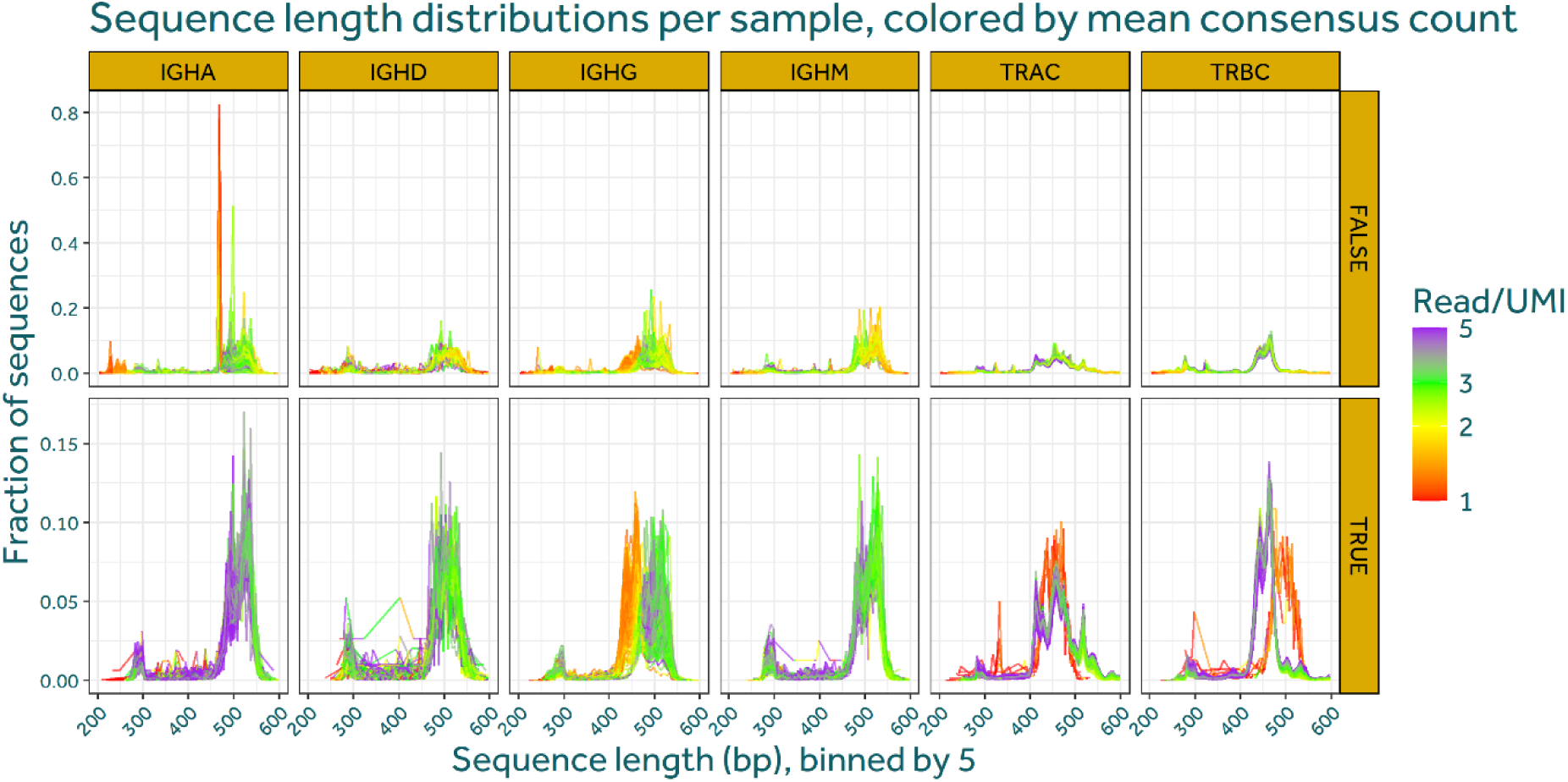
Sequence length distributions derived from collapsed UMI tables, binned in 5 bp intervals and coloured by mean reads per UMI, shown separately for all detected UMIs and those retained during assembly and filtering. A prominent IGHA off-target peak around 465 bp with low read-per-UMI support is evident in unassembled data.

**Supplementary Figure 2 –.**
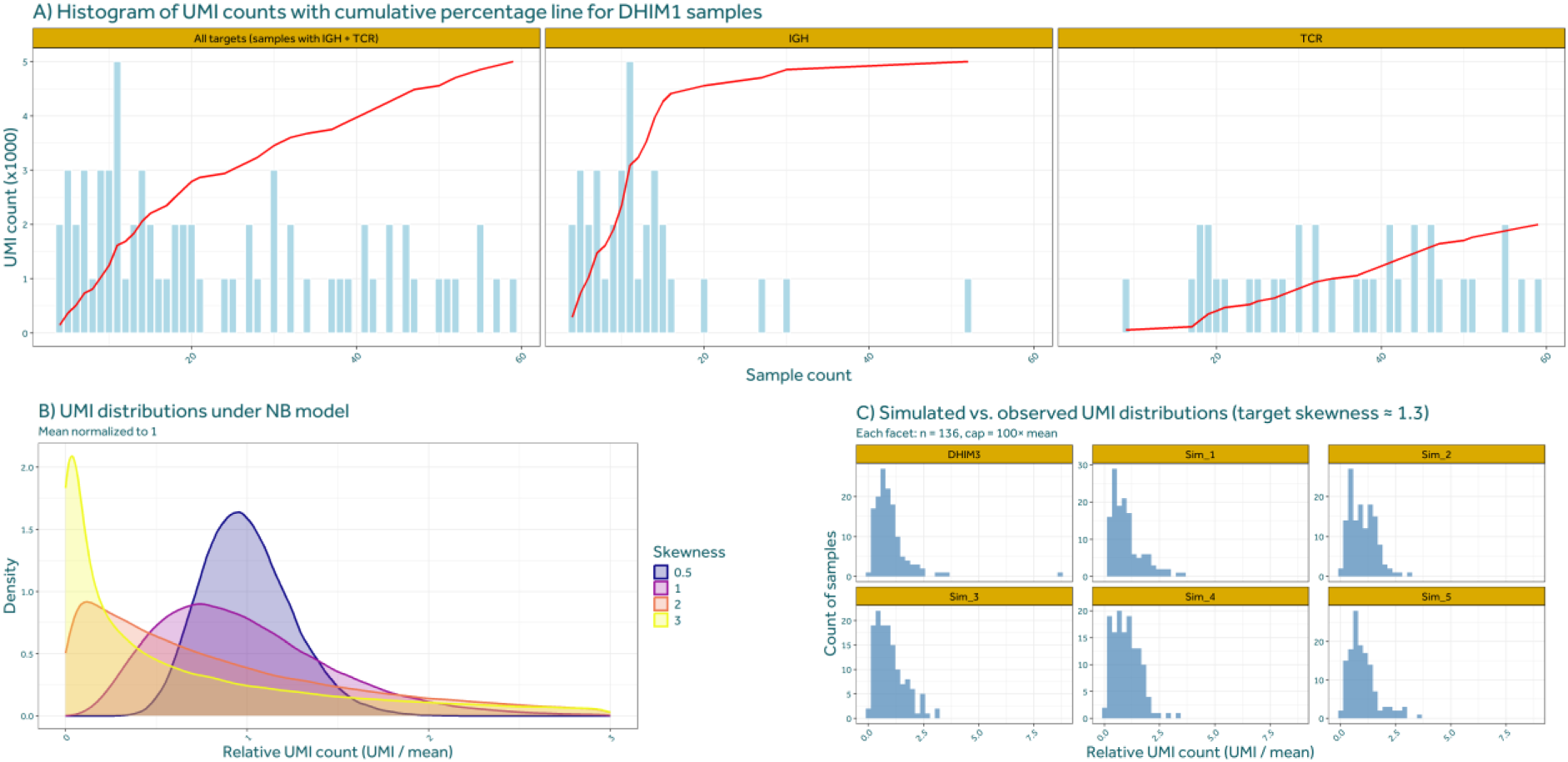
Characterization and modelling of UMI count distributions used for equi-depth pooling simulations. (A) Empirical distribution of UMI counts across DHIM1 samples, shown as histograms with an overlaid cumulative percentage curve, illustrating the strong between-sample heterogeneity in library sizes for all target sequences as well as for IGH and TCR separately. (B) Theoretical UMI abundance distributions generated under a negative binomial model with the mean normalized to one, demonstrating how increasing skewness alters the relative contribution of large versus small libraries within a pool. (C) Comparison of simulated UMI distributions with experimentally observed DHIM3 distributions for a target skewness of approximately 1.3. Each panel represents an independent simulation replicate, showing close qualitative agreement between modelled and observed relative UMI count distributions. Together, these panels motivate the choice of negative binomial models and skewness parameters used to estimate sequencing cost savings of equi-depth versus equimolar pooling.

## Notes

### Competing Interest Statement

The authors have declared no competing interest.

## Bibliography

1. Greiff, V., Miho, E., Menzel, U., and Reddy, S.T. (2015). Bioinformatic and Statistical Analysis of Adaptive Immune Repertoires. Trends Immunol. 36, 738–749. 10.1016/j.it.2015.09.006.

2. Eugster, A., Bostick, M.L., Gupta, N., Mariotti-Ferrandiz, E., Kraus, G., Meng, W., Soto, C., Trück, J., Stervbo, U., and Luning Prak, E.T. (2022). AIRR Community Guide to Planning and Performing AIRR-Seq Experiments. In Methods in Molecular Biology, pp. 261–278. 10.1007/978-1-0716-2115-8_15.

3. Chaudhary, N., and Wesemann, D.R. (2018). Analyzing immunoglobulin repertoires. Front. Immunol. 9, 1–18. 10.3389/fimmu.2018.00462.

4. Yaari, G., and Kleinstein, S.H. (2015). Practical guidelines for B-cell receptor repertoire sequencing analysis. Genome Med. 7, 1–14. 10.1186/s13073-015-0243-2.

5. Mhanna, V., Bashour, H., Lê Quý, K., Barennes, P., Rawat, P., Greiff, V., and Mariotti- Ferrandiz, E. (2024). Adaptive immune receptor repertoire analysis. Nature Reviews Methods Primers 4, 6. 10.1038/s43586-023-00284-1.

6. Robinson, W.H. (2015). Sequencing the functional antibody repertoire — diagnostic and therapeutic discovery. Nat. Rev. Rheumatol. 11, 171–182. 10.1038/nrrheum.2014.220.

7. Dahal-Koirala, S., Balaban, G., Neumann, R.S., Scheffer, L., Lundin, K.E.A., Greiff, V., Sollid, L.M., Qiao, S.W., and Sandve, G.K. (2022). TCRpower: Quantifying the detection power of T-cell receptor sequencing with a novel computational pipeline calibrated by spike-in sequences. Brief. Bioinform. 23, 1–14. 10.1093/bib/bbab566.

8. Soto, C., Bombardi, R.G., Branchizio, A., Kose, N., Matta, P., Sevy, A.M., Sinkovits, R.S., Gilchuk, P., Finn, J.A., and Crowe, J.E. (2019). High frequency of shared clonotypes in human B cell receptor repertoires. Nature 566, 398–402. 10.1038/s41586-019-0934-8.

9. Gabernet, G., Marquez, S., Bjornson, R., Peltzer, A., Meng, H., Aron, E., Lee, N.Y., Jensen, C., Ladd, D., Polster, M., et al. (2024). nf-core/airrflow: An adaptive immune receptor repertoire analysis workflow employing the Immcantation framework. PLoS Comput. Biol. 20, e1012265. 10.1371/JOURNAL.PCBI.1012265.

10. Rosenfeld, A.M., Meng, W., Chen, D.Y., Zhang, B., Granot, T., Farber, D.L., Hershberg, U., and Prak, E.T.L. (2018). Computational evaluation of B-cell clone sizes in bulk populations. Front. Immunol. 9, 338868. 10.3389/fimmu.2018.01472.

11. Wrammert, J., Onlamoon, N., Akondy, R.S., Perng, G.C., Polsrila, K., Chandele, A., Kwissa, M., Pulendran, B., Wilson, P.C., Wittawatmongkol, O., et al. (2012). Rapid and Massive Virus-Specific Plasmablast Responses during Acute Dengue Virus Infection in Humans. J. Virol. 86, 2911–2918. 10.1128/jvi.06075-11.

12. Frank, M.S., Fuß, J., Steiert, T.A., Streleckiene, G., and Gehl, J. (2021). Benchmark Quantifying sequencing error and effective sequencing depth of liquid biopsy NGS with UMI error correction. Biotechniques 70, 1–7. 10.2144/btn-2020-0124.

13. Muller, B.H., Mollon, P., Santiago-Allexant, E., Javerliat, F., and Kaneko, G. (2019). In-depth comparison of library pooling strategies for multiplexing bacterial species in NGS. Diagn. Microbiol. Infect. Dis. 95, 28–33. 10.1016/J.DIAGMICROBIO.2019.04.014.

14. Instruction manual NEBNext ® Immune Sequencing Kit (Human) version 2.0_12/21 (2021). Preprint at New England Biolabs.

15. Waickman, A.T., Lu, J.Q., Fang, H.S., Waldran, M.J., Gebo, C., Currier, J.R., Ware, L., Van Wesenbeeck, L., Verpoorten, N., Lenz, O., et al. (2022). Evolution of inflammation and immunity in a dengue virus 1 human infection model. Sci. Transl. Med. 14. 10.1126/scitranslmed.abo5019.

16. Verdonckt, T.W., Vermeersch, A.S., Struyfs, C., Nieuwerburgh, F. Van, Waickman, A., and Lagatie, O. (2025). Establishment of ‘ natural antibodies ’ during primary dengue infection. 4, 1–23.

